# A micro-evolutionary change in target binding sites as a key determinant of Ultrabithorax function in *Drosophila*

**DOI:** 10.1101/2021.08.17.456507

**Authors:** Soumen Khan, Saurabh J. Pradhan, Guillaume Giraud, Françoise Bleicher, Rachel Paul, Samir Merabet, LS Shashidhara

## Abstract

Hox genes encode Homeodomain-containing transcription factors, which specify segmental identities along the anterior-posterior axis. Functional changes in Hox genes have been directly implicated in the evolution of body plans across the metazoan lineage. The Hox protein Ultrabithorax (Ubx) is expressed and required in developing third thoracic (T3) segments in holometabolous insects studied so far, particularly, of the order Coleoptera, Lepidoptera and Diptera. Ubx function is key to specify differential development of the second (T2) and T3 thoracic segments in these insects. While Ubx is expressed in the third thoracic segment in developing larvae of Hymenopteran *Apis mellifera*, the morphological differences between T2 and T3 are subtle. To identify evolutionary changes that are behind the differential function of Ubx in these two insects, which are diverged for more than 350 million years, we performed comparative analyses of genome wide Ubx-binding sites between *Drosophila* and *Apis*. Our studies reveal that a motif with a TAAAT core is a preferred binding site for Ubx in *Drosophila*, but not in *Apis*. Biochemical and transgenic assays suggest that in *Drosophila*, TAAAT core sequence in the Ubx binding sites is required for Ubx-mediated regulation of two of its target genes studied here. *CG13222*, a gene that is normally upregulated by Ubx and *vestigial* (*<u>vg</u>*), whose expression is repressed by Ubx in T3. Interestingly, changing the TAAT site to a TAAAT site was sufficient to bring an otherwise unresponsive enhancer of the *vg* gene from *Apis* under the control of Ubx in a *Drosophila* transgenic assay. Taken together, our results suggest an evolutionary mechanism by which critical wing patterning genes might have come under the regulation of Ubx in the Dipteran lineage.

## Introduction

The diversification of morphology and emergent function of appendages have been important features of animal evolution. This is best exemplified in the diversity that is seen in closely related species of Arthropoda. In Bilaterians, first discovered in *Drosophila melanogaster, Hox* genes regulate the development of serially homologous structures along the anterior posterior axis (Carroll 1995; Lewis 1978; McGinnis and Krumlauf 1992). Subsequently, large number of studies have shown that variations in Hox gene functions, due to changes in their (i) copy number, (ii) primary structure at protein levels, (iii) their expression patterns and (iv) regulation of downstream target genes, are critical for morphological diversification.

In *Drosophila melanogaster*, the Hox protein Ultrabithorax (Ubx) specifies the development of halteres in the third thoracic (T3) segment (for detailed review see Lewis, 1978 and Khan et al., 2020). Loss of function mutations of Ubx give rise to four winged flies with complete duplication of the T2 segment in the place of T3, while overexpression of Ubx in larval wing discs leads to wing to haltere transformations (Lewis, 1978; Castelli-Gair et al., 1990; Cabrera et al., 1985; White and Akam, 1985; White and Wilcox, 1985). While Ubx is expressed in hindwing primordia of different insect species (Carroll 1995; Prasad, Tarikere et al. 2016), the outcome of its expression is not the same in all insect groups. In the Coleopteran *Tribolium castaneum* and the Lepidopteran *Precis coenia* and *Bombyx mori* Ubx is expressed only in T3 segment as in Dipteran *Drosophila*. Functional studies have shown that Ubx is required for the suppression of elytra and the specification of hindwings in the T3 of Tribolium (Tomoyasu et al. 2005), it is required to specify differences in eyespot patterns between the hindwings and forewings in *Precis* (Weatherbee et al. 1999; Matsuoka and Monteiro 2021). However, in Hymenopterans such as *Apis mellifera*, wherein T2 and T3 are distinguished by a marginally smaller hindwing, Ubx is expressed in both forewing and hindwing primordia (Prasad, Tarikere et al. 2016). The expression, however, is stronger in the hindwing primordia (Prasad, Tarikere et al. 2016). Over-expression of Ubx derived from *A. mellifera or B. mori* or *T. castaneum* can suppress wing development and specify haltere fate in transgenic *D. melanogaster* (Prasad, Tarikere et al. 2016), suggesting that changes in Ubx at the protein level may either have not contributed or marginally contributed to the evolution of its function. A study comparing genome-wide targets of Ubx in developing halteres of *D. melanogaster* and developing hindwings of *A. mellifera* and *B. mori* revealed that a large number of genes have remained common targets of Ubx in these three species (Prasad, Tarikere et al. 2016), which are diverged for nearly 350 million years. Among those common targets, a few wing patterning genes are differentially expressed in *D. melanogaster*, but not in *A. mellifera* or *B. mori* (Prasad, Tarikere et al. 2016).

To identify evolutionary changes in the regulatory sequences of targets of Ubx that may have brought some of the wing patterning genes under the regulation of Ubx, we performed comparative analyses of genome wide Ubx-binding motifs between *D. melanogaster* and *A. mellifera*. We find that while in *Drosophila*, a motif with a core TAAAT sequence (MATAAATCAY) (hereafter referred to as TAAAT motif) was enriched by Ubx in ChIP pulled-down sequences, the same was not observed in *A. mellifera*. Using a combination of in-vitro, cell culture and transgenic assays, our results reveal that the TAAAT motif is important (and bears greater relevance as compared to the TAAT motif) for the Ubx mediated-regulation of its target gene, CG13222, in *Drosophila*. We then analyzed the significance of the TAAAT motif in the Ubx mediated down-regulation of a pro-wing gene, *vestigia*l, in *Drosophila* as against the TAAT motif in its orthologue in *Apis*. In transgenic *D. melanogaster*, the enhancer of *vg* of *A. mellifera* drives reporter gene expression in both wing and halteres. We show that changing the TAAT motif to TAAAT motif was sufficient to bring this enhancer of *vg* from *Apis* under the regulation of Ubx. Taken together, our study suggests that evolutionary changes in the regulatory sequences in the targets of Ubx in dipteran lineages, such as from TAAT to TAAAT as core sequence in Ubx-binding motifs, may have been an important mechanism contributing to the evolution of wing to haltere appendages.

## Results

### Motif with a TAAAT core sequence is enriched for direct targets of Ubx in developing halteres in *D. melanogaster* but not for developing hindwings in *A. mellifera*

Previous studies indicate that despite the differences in the hindwing morphology, the Ubx protein is expressed in the third thoracic segment of both insects (as well as other insect lineages) (Fig. 1A) (Carroll 1995; Prasad, Tarikere et al. 2016). Additionally, Ubx is highly conserved across *A. mellifera, B. mori, T. castaneum, Junonia coenia* and *D. melanogaster* at the level of its DNA-binding homeodomain and the protein interaction domains, YPWM and UbdA (Fig. S1 A). We have previously shown that overexpression of Ubx from different insect species in the wing imaginal disc of transgenic *Drosophila* resulted in wing to haltere transformation in a pattern similar to *Drosophila* Ubx (Prasad, Tarikere et al. 2016). A triple mutant combination of alleles, *abx bx3* pbx, results in complete transformation of T3 to T2 (Lewis 1978). This four-wing phenotype is rescued when Ubx is over-expressed exogenously during development using a GAL4-UAS system. This phenotype can also be rescued by expressing Ubx derived from *Apis* or *Bombyx* (Merabet, S; unpublished results). This suggests that differences in insect hindwing morphology is a consequence of events occurring downstream of or in parallel to Ubx. Comparison of targets of Ubx between *D. melanogaster*, and *A. mellifera* suggest that many functionally important genes are common to the two lineages, although many key genes are differentially expressed between T2 and T3 in *Drosophila*, but in *Apis* (Prasad, Tarikere et al., 2016; Fig. S1 B). This reconfirms that morphological differences may lie at the levels of regulation of those targets.

**Figure 1:**
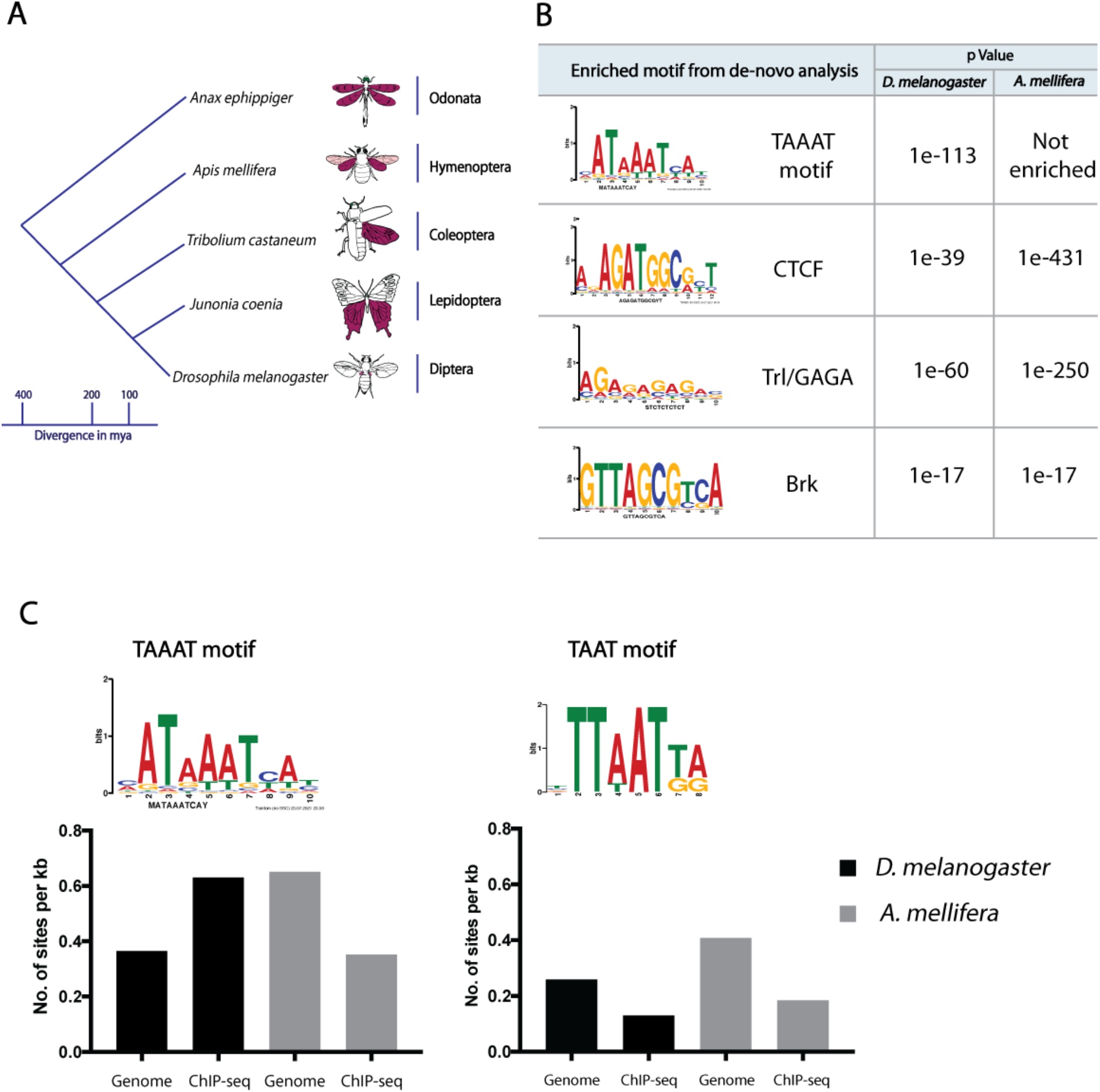
Motif with a TAAAT core sequence is enriched in direct targets of Ubx in developing halteres in *D. melanogaster* but not in developing hindwings in *A. mellifera*. **A)** While Ubx is required for haltere development in Dipterans such as *Drosophila melanogaster*, differences in hindwing morphology in the third thoracic segment of diverse insects is not correlated to the mere presence of Ubx. **B)** De-novo motif analysis of Ubx ChIP-seq in *Apis* hindwings and *Drosophila* halteres reveal enrichment of the TAAAT motif specifically in *Drosophila* but not in *Apis*. Binding sites for transcription factors like Trl, CTCF and Brk are, however, enriched in both datasets. **C)** FIMO analysis of the TAAAT motif reveals a 1.7fold enrichment of sites in *Drosophila* ChIP-seq sequences as compared to the entire genome. No such enrichment was found for the *Apis* ChIP-seq datasets. The TAAT motif was not enriched in either of the datasets.

We carried out ChIP-Seq for Ubx from the third instar haltere discs (GSE205177). This method provided better resolution over earlier methods used by us (Agrawal et al., 2011) and others (Choo et al., 2011; Slattery et al., 2011) to identify Ubx-specific binding motifs. We then performed a comparative analysis of Ubx ChIP-Seq data generated from *D. melanogaster* haltere imaginal discs to Ubx ChIP-seq data generated from *A. mellifera* hindwing discs (reported in Prasad Tarikere et al., 2016).

De-novo motif analysis of ChIP-Seq data from *D. melanogaster* showed significant enrichment for a motif with a TAAAT core sequence (MATAAATCAY) (henceforth referred to as the TAAAT motif) as putative Ubx-binding site (Fig 1B). We observed that enrichment for TAAAT motif is only in *Drosophila* and not in *Apis*, while both the ChIP-Seq datasets showed enrichment for binding sites for other transcription factors such as GAGA, CTCF and Brk (Fig. 1B). We calculated the frequency of the TAAAT motif in the *Drosophila* dataset and found it to be 1.7X in Ubx-pulled down sequences as compared to its prevalence in the whole genome. Interestingly, the frequency of the TAAAT motif in the entire genome was 1.8 fold higher in *A. mellifera* as compared to *D. melanogaster* (Fig. 1C), suggesting that the TAAAT motif is selectively enriched in the Ubx bound regions only in *D. melanogaster* and not in A. *mellifera*.

We also observe, similar to previous results (Agrawal et al. 2011), that the canonical TAAT motif is not enriched in Ubx-pulled down sequences, rather it was found under-represented compared to its prevalence in the whole genome (Fig. 1C). This was found to be true for both *Drosophila* and *Apis* datasets.

### The TAAAT motif is critical for Ubx mediated regulation of a target gene in *Drosophila* halteres

Our results suggest that the TAAAT motif is enriched in Ubx-pulled down sequences from haltere imaginal discs of *Drosophila*, but not from developing hindwings of *A mellifera*. Earlier studies have reported that the TAAAT motif is bound by the Ubx-Extradenticle-Homothorax complex (Slattery et al. 2011; Sánchez-Higueras et al. 2019). However, neither Extradenticle (Exd) nor Homothorax (Hth) are expressed or required for the development of the haltere capitellum (Casares and Mann 2000). In this context, we sought to determine the functional significance of Ubx-binding to the TAAAT motif during haltere development.

We first compared Ubx ChIP-seq data from *Drosophila* halteres to RNA seq data (GSE205352) for *Drosophila* wing and haltere imaginal discs to identify those direct targets that are upregulated or downregulated in the haltere. We then compared the frequency of the TAAAT motif in Ubx bound response elements of putative direct targets that are upregulated or downregulated between wing and haltere discs. We found the frequency of the TAAAT motif in Ubx bound response elements in the upregulated category is marginally higher (1.4 times) as compared to the downregulated category (Table S2). This suggests that while the TAAAT motif may be preferred site for Ubx to bind on the enhancers of its targets, Ubx may use this motif to regulate both upregulated and downregulated targets in the haltere.

We next evaluated the affinity and functional importance of Ubx binding to the TAAAT motif in one of the well characterized targets, *CG13222*, which is upregulated by Ubx in developing halteres (Mohit et al. 2006; Hersh et al. 2007). The “*edge*” enhancer of *CG13222* has two TAAT motifs at 17 bp apart (CG13222_WT, Fig 2A). Hersh et al. (2007) have named the two motifs as site 1 and site 2. In their transgenic assays, site1 appears to be critical for Ubx-mediated regulation of *CG13222*, while site2 is dispensable. Interestingly, the site1 in the *edge* enhancer, which is critical for its regulation by Ubx, has a TAAAT motif which overlaps with the TAAT motif (Fig. 2A). Their analysis of site1 involved mutating both TAAT and TAAAT motifs, which leads to loss of enhancer driven reporter gene expression (Hersh et al. 2007). We mutated site1 in such a way that the mutant enhancers carry either the TAAAT motif (CG13222_M1_A) or the TAAT motif (CG13222_M1_B) (Fig. 2A).

**Figure 2:**
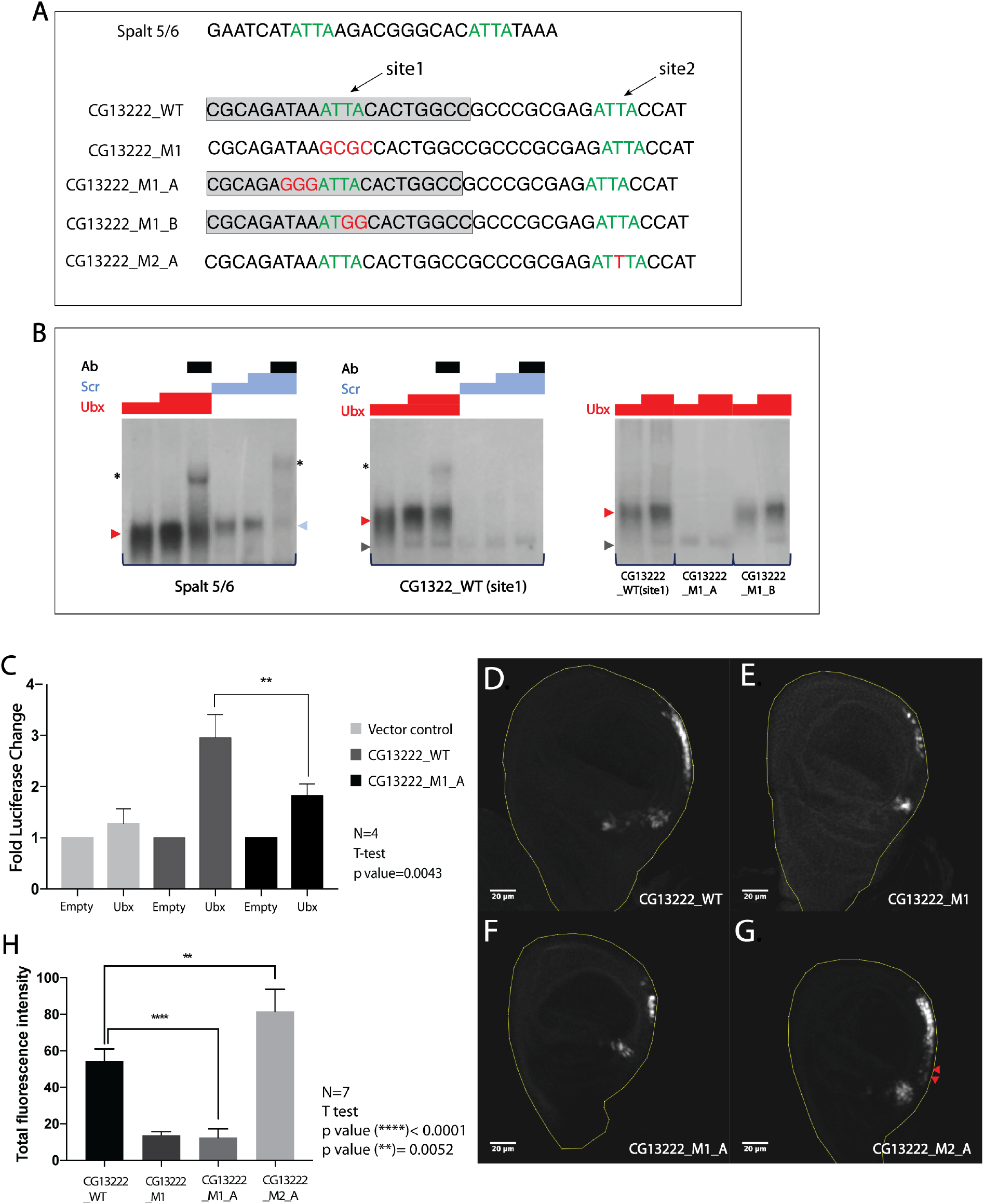
The TAAAT motif is critical for Ubx mediated regulation of a target gene in *Drosophila* halteres. **A)** Sequence of part of *sal* enhancer (Spalt 5/6), *CG13222* enhancer and mutations of the *CG13222* enhancer (CG13222_M1_A, CG13222_M1_B) used as probes for EMSA. The wild type *CG13222* enhancer has two Ubx binding sites (shown in green) termed here as CG13222_WT (site1) and CG13222_WT (site2). Mutations were generated to specifically modify either the TAAT (CG13222_M1_B, CG13222_M2_A) or the TAAAT (CG13222_M1_A) motifs while keeping the other one intact in site1 of the *CG13222* enhancer. For Luciferase and Transgenic assays, the entire 512bp spanning the *CG13222* edge enhancer was used. **B)** While both Ubx and Scr bind to TAAT motif of *sal*, only Ubx binds to TAAAT motif of CG13222_WT (site1). Binding of Ubx to CG13222_WT (site1) is severely reduced when the TAAAT motif is mutated to TAAT motif (CG13222_M1_A), whereas no such effect is seen on mutating the TAAT motif (CG13222_M1_B). **C)** The *CG13222* enhancer drives Luciferase activity in a S2 cell culture system in presence of Ubx. Significant loss of reporter activity is observed in the mutant enhancer (CG13222_M1_A) where the TAAAT motif is affected. For statistical analysis, t-test was performed (two-tailed). **D-H)**. The *CG13222* enhancer drives reporter expression in the posterior edge of the third instar haltere imaginal discs (D). The *CG13222_M1* mutant where both the TAAT and TAAAT motifs are affected show a marked reduction in reporter expression (E). The *CG13222_M1_A* mutant carrying changes in the TAAAT site of the enhancer has a similar phenotype as the *CG13222_M1* mutant (F) and shows a significant reduction in GFP expression as seen from calculation of average fluorescence intensity (H). In the *CG13222_M2_A* enhancer where the TAAT is mutated to TAAAT in site2 results in significantly higher level of reporter expression (H) along with ectopic expression in some domains (shown by red arrows) (G). For statistical analysis, t-test was performed (two-tailed).

To examine the relative affinities of Ubx binding to the TAAAT and TAAT sites by Electrophoretic Mobility Shift Assay (EMSA), we used a 21bp long probe covering site 1 of *edge* enhancer of *CG13222* (nucleotide stretch in the grey box in Fig. 2A). We find that Ubx can bind to the wild type probe derived from the *CG13222 edge* enhancer. Mutations in the TAAAT motif in site1 of *CG13222* lead to a substantial loss of this binding (probes CG13222_M1_A, Fig. 2B). Conversely, mutating the TAAT motif alone had no significant effects on binding of Ubx to the probe (CG13222_M1_B, Fig S2 A).

We also observed that another Hox protein, Scr, was unable to bind to the wildtype probe *CG13222*. On the other hand, both Ubx and Scr bind to a 29bp probe derived from the *sal* enhancer (Galant et al. 2002) containing TAAT sites (Fig 2B). Perhaps, the TAAAT motif provides higher specificity of binding for Ubx as compared to other Hox proteins.

Next we examined the importance of the TAAAT motif in gene regulation using Luciferase reporter assays in S2 cells and haltere imaginal discs in transgenic *Drosophila*. For these functional assays, we used a 512bp long region spanning the *edge* enhancer of *CG13222* as reported by Hersh et al. (2007), which has both site1 and site2. Consistent with *CG13222* being a target of Ubx upregulated during haltere development, in the presence of Ubx a luciferase expression construct driven by the enhancer of the gene was significantly upregulated (2.8 folds) in S2 cells (Fig. 2C). Using a similar experimental design, we observed a significant decrease in Ubx-dependent expression when TAAAT alone is mutated (construct CG13222_M1_A in Fig. 2C) compared to the wildtype *CG13222* enhancer.

We next generated transgenic strains of *D. melanogaster* carrying the wild type or mutant enhancers (as explained above) cloned upstream of a GFP reporter. The wildtype *CG13222* enhancer showed GFP expression at the posterior edge of the haltere imaginal discs consistent with earlier reports (Fig. 2D) (Mohit et al. 2006; Hersh et al. 2007). We first replicated the mutation described in Hersh et al. 2007 wherein both the TAAT and TAAAT motifs are mutated (CG13222_M1) and found that the reporter gene expression is significantly reduced when driven by the mutant enhancer (Fig. 2E). We then analysed the CG13222_M1_A mutant where the TAAAT motif is specifically mutated while the TAAT motif is maintained intact and observed a significant loss of reporter expression (Fig. 2F, 2H). However, the pattern of GFP expression patterns driven by CG13222_M1 and CG13222_M1_A enhancers were similar, suggesting that the TAAAT motif is specifically required for Ubx-mediated activation of the *CG13222* enhancer in *Drosophila* halteres.

As described earlier, the *edge* enhancer of *CG13222* has a second TAAT motif (site2 in Fig 2A), which is redundant for Ubx-mediated upregulation of *CG13222* in the haltere discs (Hersh et al. 2007). We also attempted to examine the effect of mutating the TAAT motif to a TAAAT motif (in site2, construct CG13222_M2_A), which results in two high-affinity binding motifs in the *edge* enhancer of *CG13222*. We observed a significant reduction in the luciferase activity (Fig. S2 B), perhaps, due to negative effect of two high-affinity binding sites for Ubx in close proximity. However, the *Drosophila* transgenic line carrying the same mutation showed significantly stronger expression of the GFP reporter (Fig 2G, 2H) compared to the wildtype *CG13222* enhancer. Enhancer driven expression was also observed in ectopic regions in the posterior compartment (red arrows in Fig. 2G), suggesting that the presence of an additional TAAAT motif may bring the otherwise unresponsive site2 under Ubx regulation.

### The TAAAT motif causes Ubx mediated repression of an orthologous enhancer of the *vestigial* gene from *A. mellifera* in a transgenic *Drosophila* assay

We had previously reported that several wing patterning genes such as the *vestigial* (*vg*) gene are expressed in both fore- and hindwings primordia in *A. mellifera*. On the contrary, in *Drosophila, vg* is a direct target of Ubx (Suppl. 2C) and is downregulated in the developing halteres (Galant et al. 2002; Hersh and Carroll 2005). Of the two well characterized enhancers, the quadrant enhancer of *vg* (henceforth named as *quad-vg*) is differentially expressed between the wing and haltere imaginal discs. In a transgenic *Drosophila* assay, an enhancer of *vg* from *Apis* (henceforth named as *Apis-vg*) equivalent to *quad-vg* showed similar patterns and levels of expression between wing and haltere discs (Prasad, Tariekre et al. 2016), suggesting that *Drosophila* Ubx too cannot repress the *Apis-vg* enhancer. We evaluated the role of TAAAT in *Drosophila* as against TAAT in *Apis* in the regulation of the *quad-vg* and *Apis-vg* enhancers in transgenic assays.

We first scanned the entire 850bp sequence of the *quad-vg* enhancer and the 575bp sequence of the *Apis-vg* enhancer for Ubx binding motifs, TAAAT and TAAT. We observed a 25bp cassette containing both TAAT and TAAAT motifs in the *quad-vg* enhancer (Fig 3A). In the Apis-vg enhancer, we found only a single TAAT motif and did not find any TAAAT motif (Fig 3A). In an attempt to test the significance of this difference in Ubx binding motifs between the two enhancers, we generated several *Drosophila* transgenics carrying the wild-type and mutated *quad-vg* and *Apis-vg* constructs (mutations in TAAT or in TAAAT motifs) upstream of a GFP reporter (Fig 3A, Suppl. Fig 2D, Suppl. Fig 3A). As reported earlier, we observed that the wild type *quad-vg* enhancer drives expression of the GFP reporter in the wing imaginal discs, albeit at much lower levels (compared to earlier reported *quad-vg -*lacZ reporter), but not in the haltere imaginal discs (Suppl. Fig 2E). However, owing to complete loss of enhancer readout in both wing and haltere imaginal discs in the transgenic carrying mutations in the TAAAT motif of the *quad-vg* enhancer (Suppl. Fig 2E), we were unable to conclude the role of the TAAAT motif or the TAAT motif in Ubx mediated regulation of the *quad-vg* enhancer.

**Figure 3:**
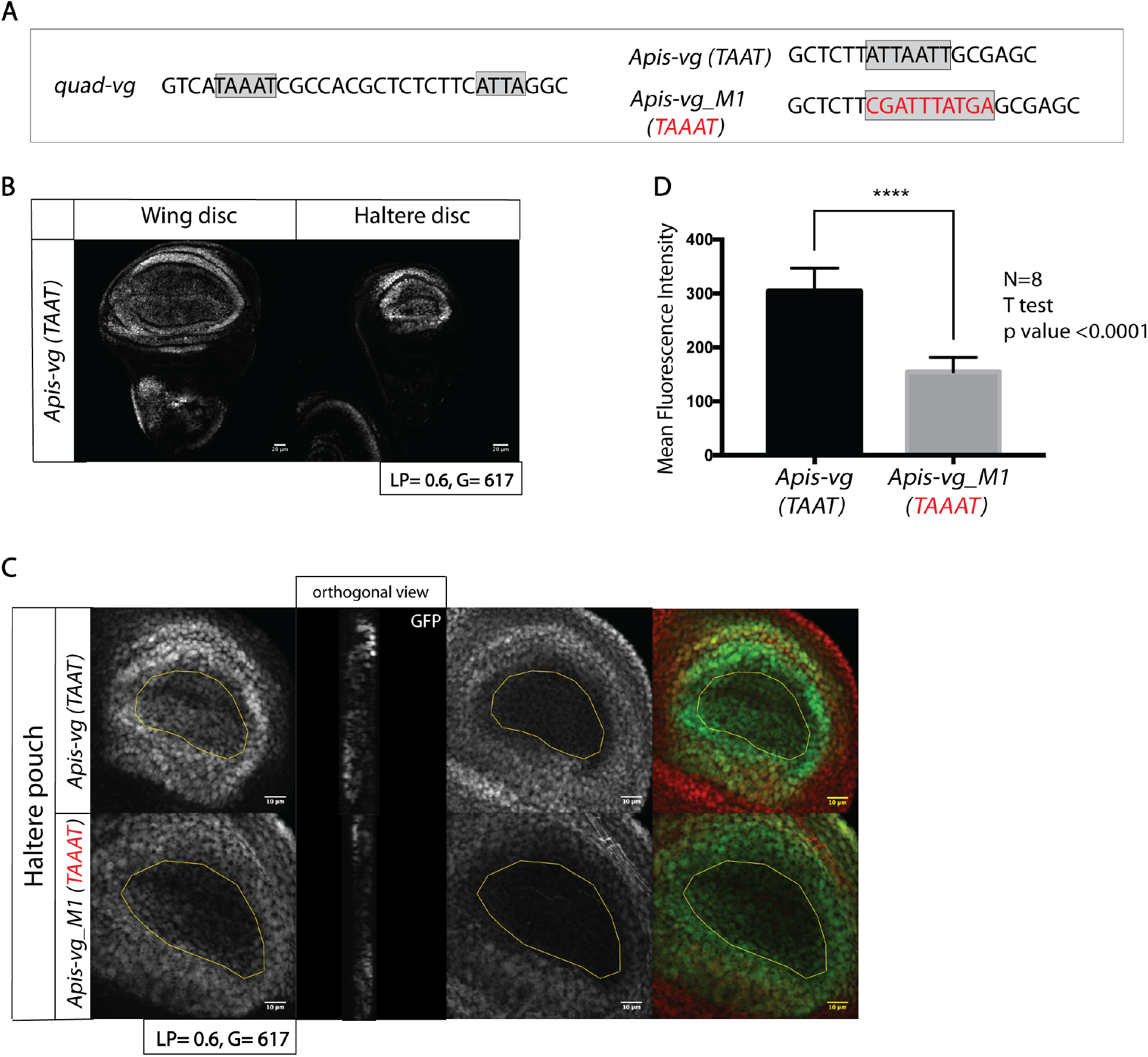
The TAAAT motif causes Ubx mediated repression of an orthologous enhancer of the *vestigial* gene from *A. mellifera* in a transgenic *Drosophila* assay. **A)** Sequence of part of the 805bp enhancer of the *vestigial* gene in *Drosophila (quad-vg)* containing both TAAT and TAAAT motifs and part of the 575bp enhancer of the *vestigial* gene in *Apis (Apis-vg)*, showing the presence of a TAAT motif. **B)** The Apis-vg enhancer drives similar expression of reporter GFP in hinge and pouch regions of wing and haltere imaginal discs. **C)** The mutant enhancer (*Apis-vg_M1*) with introduced TAAAT motif shows reduced levels of GFP expression in the haltere pouch. Magnified images of the haltere pouch of *Drosophila* transgenics expressing GFP under *Apis-vg* and *Apis-vg_M1* enhancers. Note much reduced GFP levels specifically in the haltere pouch of *Apis-vg_M1* transgene. Orthogonal views of the haltere imaginal pouch indicate a clear difference in GFP expression driven by the WT and the mutant enhancers. Hth staining, which is hinge-specific is used to demarcate the pouch region. **D)** Quantification of average fluorescence intensity in the haltere pouch for the *Apis-vg* and *Apis-vg_M1* transgenics. For statistical analysis, t-test was performed (two-tailed).

We next analysed the reporter GFP expression in wing and haltere imaginal discs of *Drosophila* transgenics carrying the wild type and mutated *Apis-vg* enhancers. As reported earlier (Prasad, Tarikere et al., 2016), we observed similar expression of GFP in both wing and haltere imaginal discs (Fig. 3B). Interestingly, in a chimeric enhancer (*Apis-vg_M2*), where the 25bp Ubx-binding cassette from the *Drosophila quad-vg* enhancer replaced its counterpart in the *Apis-vg* enhancer (Suppl. Fig. 3A (iii)), we observed differential expression of the GFP reporter between wing and haltere imaginal discs (Suppl. Fig. 3B). Its expression was much lower in the haltere pouch compared to the GFP reporter driven by the wildtype *Apis-vg* enhancer. The loss of reporter GFP expression in the pouch region was consistent across all mutant *Apis-vg* constructs which bore a TAAAT motif but was not observed in constructs where the TAAAT motif was absent (Suppl Fig 3A (vi), 3B). More importantly, changing only the TAAT motif of the *Apis-vg* enhancer to a TAAAT motif (*Apis-vg_M1*), was sufficient for the repression of the GFP reporter in the pouch (Fig. 3C, 3D,Suppl. Fig 3B) of haltere discs without affecting its expression in the wing pouch (Suppl. Fig 3C).

Taken together, our results reveal that a microevolutionary change in the enhancer sequence, specifically modifying the TAAT motif to a TAAAT motif, may have brought certain critical wing-patterning genes, such as the pro-wing gene *vg*, under the regulation of Ubx during the evolution of dipterans.

## Discussion

Genetics, cell and molecular biology methods have established, unequivocally, the role of Hox genes as master control genes in regulating segment-specific developmental pathways and diversification of body plans during evolution. However, precise mechanism by which orthologous Hox proteins mediate differential development of a specific morphological feature in different species is still largely unknown. Main reasons this has eluded systematic study include their highly conserved DNA-binding domains across all species.

In *Drosophila*, the Ubx protein is expressed in the T3 segment and specifies the development of the haltere by activating or repressing a number of genes at various hierarchical levels of wing patterning (Weatherbee et al. 1998; Shashidhara et al. 1999). Ubx is also expressed in the T3 segment of other insect species and thought to specify different fates in each of them. For example, in Coleopterans, Ubx specifies the development of hindwings instead of elytra and in Lepidopterans it specifies differences in eyespot patterns between the forewing and hindwings. In more ancestral Hymenopterans such as *Apis mellifera*, which have two pairs of wings, hindwings are marginally smaller than the forewings. In A. mellifera, Ubx is expressed in both forewing and hindwing primordia, although its expression is stronger in T3 (Prasad, Tarikere et al. 2016). It is possible that divergence of Coleopterans, Lepidopterans and Dipterans involve both suppression of Ubx expression in T2 and Ubx acquiring ability to regulate various developmental pathways in T3.

To identify the mechanisms governing the differential regulation of wing patterning genes between T2 and T3 in *Drosophila*, we employed a comparative genomics approach and identified that the TAAAT motif is enriched by Ubx specifically in the enhancers of targets of Ubx in *Drosophila* halteres but not in *Apis* hindwings. Our functional studies reported here suggest that Ubx binds to the TAAAT motif with higher affinity than to the TAAT motif and its binding to former is critical for the up-regulation of *CG13222* expression in the haltere imaginal discs.

An earlier report of ours (Prasad, Tarikere et al. 2016) and this report suggest that unlike the *quad-vg* enhancer of *Drosophila*, an enhancer of *vg* gene of *A. mellifera* is not differentially expressed, but drives GFP expression in both wing and haltere imaginal discs in a transgenic assay. We show that changing the TAAT motif to TAAAT motif, dramatically, brought the enhancer under the negative regulation of Ubx. The *Apis-vg* enhancer, thus, allowed us to dissect the importance of TAAAT motif in dipteran evolution. As *vg* is a pro-wing selector gene and can assign wing fate to any group of dorsal epithelial cells (Kim et al. 1996; Klein and Arias 1998; Williams et al. 1991; Neumann and Cohen 1996), it is a critical target of Ubx to specify haltere development. In this context, a microevolutionary change (TAAT to TAAAT motif) in the enhancer of an ancestral *vg* may be a critical step in the evolution of dipterans.

It is, however, very unlikely that a large number of convergent mutations would be involved in this process, wherein multiple TAAT motifs are evolved to TAAAT motifs. A few critical genes, such as *vg*, upstream of wing patterning pathways may have evolved to become targets of Ubx through this route. Once regulatory networks of those genes are modulated by Ubx, chromatin landscape of many other targets (Ubx may bind to these targets via TAAT motifs) would change making them amenable for Ubx-mediated regulation and thereby specifying the haltere fate. In addition, certain co-factors influencing Ubx-mediated regulation of its targets irrespective of Ubx binds to TAAAT or TAAT motif cannot be ruled.

## Materials and Methods

### 1. ChIP sequencing

Third instar wandering larvae were cut, inverted and fixed with 1.5% PFA (ThermoFisher scientific) for 20 minutes at room temperature and subsequently quenched (125mM Glycine solution, 1XPBS). Samples were washed twice with 1XPBS (Sigma) for 10minutes each and dissected for wing and haltere discs (500 number of wing discs and 1000 number of haltere discs were used per replicate). Samples were lysed with Cell Lysis Buffer (10mM Tris-Cl pH 8, 10mM NaCl, 0.5% NP40, 1xPIC (Roche)) followed by mechanical shearing for 10minutes on ice. After centrifugation at 2000rpm for 10mins at 4 oC, the was pellet resuspended in Sonication buffer (50mM Tris-Cl pH 8, 1% SDS, 1% NP40, 10mM EDTA, 1xPIC) followed by incubation on ice for 30 minutes. Sonication was carried out using the Covaris S2 sonicator using settings (DC 20%, Intensity 5, Cycles of Burst 200, Time= 20 minutes). Samples were centrifuged at 14000rpm for 15mins at 4 oC and precleared using with 4ml of Magnetic A Beads (Diagenode) at 4 oC for 1 hour. Preclearing beads were removed and 2ml of Polyclonal anti-Ubx antibodies (Agrawal et al. 2011) were added with overnight incubation at 4 oC. Magnetic A beads were added and incubated for 4 hours at 4 oC. The supernatant was removed and beads washed twice with low salt buffer (20mM Tris-Cl pH 8, 150mM NaCl, 0.1 % SDS, 2mM EDTA, 1% TritonX-100), twice with High Salt buffer (20mM Tris-Cl pH 8, 500mM NaCl, 0.1 % SDS, 2mM EDTA, 1% TritonX-100), twice with LiCl buffer (10mM Tris-Cl pH 8, 250mM LiCl, 1% NP40, 1% Na-Deoxycholate, 500mM EDTA,), and once with TE buffer (10mM Tris-Cl pH 8, 500mM EDTA). Samples were eluted in elution buffer (50mM Tris-Cl, 1mM EDTA, 1% SDS, 50mM NaHCO3). Samples were decrosslinked followed by treatment with RNAse treatment at 37 degrees for 1hour and Proteinase K treatment at 42 oC for 2 hours. DNA was purified using PCI purification and quantified using the Qubit HS DNA quantification system.

For library preparation, equal amount of DNA (∼2 ng) was used as an input for NEB Ultra II DNA library prep kits (NEB #E7645). Number of cycles for amplification of adapter ligated libraries were estimated by the qPCR before final amplification to avoid any bias arising due to PCR amplification and indexing (NEB #E7350). Final amplified libraries were purified twice, first with 1X followed by 0.8x volume of beads per sample using HiPrep PCR clean up system (Magbio #AC-60050). Library concentration was determined using Qubit HS DNA kit (Invitrogen #E7350) and average fragment size was estimated using DNA HS assay on bioanalyzer 2100 (Agilent #5067-4626) before pooling libraries in equimolar ratio. Sequencing reads (100bp PE) were obtained on the Hiseq 2500 V4 platform at Macrogen Inc, Korea.

### 2. ChIP sequencing analysis

For *D. melanogaster*, raw reads were trimmed using the Trimmomatic software (command) and aligned to the dm6 genome (version BDGP6.28). Peak calling was performed using MACS2 (maxdup=1, FDR 0.01) (Zhang et al. 2008) and high confidence peaks (occurring in at least two biological replicates) were identified (https://ro-che.info/articles/2018-07-11-chip-seq-consensus). For *Apis mellifera*, raw fastq reads were downloaded from NCBI (GSE71847). Reads were aligned to the Amel_4.5 genome (version 4.5.47) and peak calling done using MACS2 (maxdup=1, FDR 0.01). High confidence peaks occurring in both replicates were identified. Peaks were annotated to their nearest TSS using the Homer software (annotatePeaks.pl) (Heinz et al. 2010). Gene ontology of targets was performed using GO (Ashburner et al. 2000; Carbon et al. 2021) and overrepresented GO terms plotted. The ChIPseeker program (Yu et al. 2015) was used to annotate peaks to genomic regions. Genome tracks were made using Deeptools (normalizeUsing RPGC, binsize 1, minMappingQuality 10, smoothLength 150) (Ramírez et al. 2016) and visualized using the IGV software (Robinson et al. 2011). Heatmaps were generated using Deeptools (referencePoint TSS).

### 3. RNA seq data generation and analysis

Third instar wandering larvae of the Canton S strain were collected, cut, inverted and dissected for wing and haltere discs in cold PBS. Experiments were carried out in triplicates and samples were snap frozen in Trizol. Library preparation and sequencing on Illumina Hiseq2000 was performed at Genotypic Technology, Bangalore, India. Raw fastq files were obtained and used for analysis. Raw reads were trimmed using Trimmomatic and aligned to the reference genome using the Hisat2 software (Kim et al. 2015, 2) followed by read counts using the Htseq software (Anders et al. 2015). Differentially expressed genes were identified using the edgeR software (Robinson et al. 2010) using 1.5 fold difference as the cut-off.

### 4. Motif analyses and protein sequence comparison

De-novo motif analysis was carried out using Homer (findMotifsGenome.pl). The PWM for the TAAAT motif obtained from Homer was converted to Transfac format using RSAT (Nguyen et al. 2018) and finally converted to MEME format using transfac2meme command. Identification of binding motifs in enhancers of *CG1322*2, *quad-Vg* and *ApisVg* was performed using the MAST software from MEME suite (Bailey et al. 2009).

Protein sequences of Ubx from different insect species were downloaded from Uniprot and multiple sequence alignment performed using Clustal Omega (Madeira et al. 2019). Alignments were visualised and processed using the Jalview software (Waterhouse et al. 2009).

### 5. Frequency calculation of TAAAT motif in Ubx response elements

Using a 1.5 fold difference (FD) cut-off, genes that are upregulated or downregulated in the haltere imaginal discs were identified. Ubx targets obtained from ChIP-seq were mapped to differentially expressed genes to identify putative Ubx response elements that are upregulated, downregulated or not-differentially expressed in the halteres. The PWM of the TAAAT motif was generated and its frequency calculated in each group of Ubx response elements using the FIMO software.

### 6. Plasmids constructs, cloning and transgenic fly generation

The full list of primers as well as sequences of enhancers of the *CG13222, quad-Vg* and *ApisVg* is provided as supplementary data (File S2). For Luciferase assays, site-directed mutagenesis was used to clone enhancer constructs of the *CG13222* gene between Kpn1 and Nhe1 restriction sites, upstream of a modified pGL3 vector containing a 5X Dorsal binding site. Metallothionein inducible pRMHa3 vectors containing Ubx were previously generated in the lab and empty pRMHa3 vector was generated by excising the cloned Ubx sequence. For generating transgenics, all enhancer constructs were cloned between the Nhe1 and Kpn1 restriction sites of the pH-stinger-attb (a kind gift from Manfred Frasch) and injected into the attp40 site on the second chromosome using a mini white screen at NCBS fly facility, Bangalore. All constructs were sequence verified before use in functional assays. Insertion of various transgenes in attp40 site helped for better comparison of wildtype and mutant transgenes of a given enhancer.

### 7. Luciferase assays

S2 cells were checked for contamination before performing Luciferase assays. Cells were plated onto 24 well plates at a density of 3*10^5 cells per well 6 hours prior to transfection. For every construct, either wild type or mutant, two sets of experiments were designed; one well was co-transfected with the enhancer construct in pGL3 vector and the empty pRMHa3 vector whereas the other well was co-transfected with the enhancer construct and pRMHa3 vector containing Ubx. Renilla luciferase was used as an internal control and co-transfected in all experiments. Transfection was carried out using the Effectene transfection reagent and all experiments carried out in 3 technical replicates and at least 3 biological replicates. 48 hours post transfection, sterile CuSO4 solution was used to induce expression of Ubx at a final concentration of 500um and incubated for 24 hours. Cells were harvested, pelleted down (1000rpm for 4mins) and 100ul of 1X Passive Lysis buffer added and vortexed to dissolve the cell pellet. Cells were incubated for 15mins at room temperature and further spun at 10,000rpm for 90secs to collect the supernatant. The luminescence was measured with the Dual-glo Luciferase assay kit (Promega) on Ensight Plate reader (Perkin Elmer). All readings were normalized to Renilla luminescence and datasets compared using the Prism software.

### 8. EMSA

VC-Ubx and VC-Scr were cloned in the PcDNA3 vector and produced with the TNT-T7-coupled in vitro transcription/translation system (Promega) for EMSAs, as previously described (Hudry et al. 2012). Shortly, between 3 and 6 microliters of programmed lysate was used for each protein (100 ng/ml of proteins were produced on average). The VC-Ubx and VC-Scr fusion proteins were produced separately (0,5mg of each plasmid was used for the in vitro transcription/translation reaction). Supershift against the VC fragment was performed by adding the anti-GFP antibody after 15 minutes in the binding reaction. Each band shift experiment was repeated at least two times.

### 9. Immunostaining and Microscopy

Wandering third instar larvae were cut and inverted in cold PBS followed by fixation with 4% PFA for 20mins with gentle rocking. Samples were washed thrice with 0.1% PBTX (0.1% Triton in PBS) for 10 minutes each, followed by one hour of blocking at room temperature (blocking solution: 0.5% BSA in 0.1%PBTX). Samples were incubated with primary antibodies (dilutions made in blocking solution) at 4 oC overnight followed by washing with 0.1% PBTX for 10mins (x3) at room temperature. Samples were incubated with secondary antibodies for one hour at room temperature, washed with 0.1% PBTX for 10mins (x3) and finally with PBS before mounting. Wing and haltere imaginal discs were dissected out, mounted using Prolong Gold Antifade (Invitrogen) and stored at 4 degrees. Antibodies used in the study are Rb-GFP (1:1000) (Invitrogen), Rb-Ubx (1:1000) (Agrawal et al., 2011), m-Ubx (1:30) (DSHB), goat-Hth (1:100) (Santa Cruz Biotechnology), Alexa Fluor 568 (1:1000) (Invitrogen) and Alexa Fluor 633 (1:1000) (Invitrogen). All images were taken on Leica Sp8 system at Microscopy Facility, IISER Pune. Laser settings (Laser Power (LP) and Gain (G)) used for measuring GFP fluorescence have been indicated for each image panel. All images were processed using the ImageJ software and compiled using Microsoft PowerPoint and Adobe Illustrator.

## Data availability

Raw fastq files of ChIP sequencing for the Ubx protein in *Apis mellifera* hindwings can be accessed from NCBI (GSE71847). Raw fastq files of ChIP sequencing for the Ubx protein in Drosophila melanogaster halteres and RNA sequencing data for wing and haltere imaginal discs can be accessed from NCBI (GSE205177 and GSE205352).

## Acknowledgments

We thank G. Deshpande and members of the LSS and SM laboratories for critical input. We thank S. Galande for providing reagents for Library preparation. This work was supported primarily by an Indo-French Research grant from CEFIPRA to SM; a JC Bose Fellowship and grant from the Department of Science and Technology, Government of India to LSS; and a Council of Scientific & Industrial Research (CSIR) Fellowship to SK.

## Author contributions

SK carried out all fly experiments, image analyses, and wrote the MS.

SK and SP did ChIP-Seq and RNA-Seq and analyses.

GG, FB and RP did EMSA experiments.

LSS and SM conceived the project and wrote the MS.

We declare “no-conflict-of-interest”.

## Notes

### Competing Interest Statement

The authors have declared no competing interest.

### Summary of Updates

The text has been modified to include some results that have been added to bolster the finding in the paper. The materials and methods section has also been updated

