## Supplementary Information for "A micro-evolutionary change in target binding sites as a key determinant of Ultrabithorax function in *Drosophila*"

**Supplementary Information, Table and Figures**

**1. Primer sequences used for cloning CG13222 constructs into pGL3-DE5 or pH-Stinger-attb** **vectors**

CG13222_FP:	ACGGTACCCATAGACCACCAGCCACTGT

CG13222_RP:	ATCGCTAGCTCAATCGCGTACCGAAGCAA

**2. Primer sequences used for cloning quad-vg constructs into pH-Stinger-attb vectors**

quad-vg_FP:	ACGGTACCGGAGCTCCCTCCGGAGAC

quad-vg_RP:	ATCGCTAGCCGATTGTACTTTGTCGTTTCTAA

**3. Primer sequences used for cloning Apis-vg constructs into pH-Stinger-attb vectors**

Apis-vg_FP:	ACGGTACCCTTCTCGCGAGAAACGAGAGGC

Apis-vg_RP:	ATCGCTAGCGTGGACAGTGACGAGGACACG

**4. Reverse Primer (for all constructs):**

GL primer2 (CTTTATGTTTTTGGCGTCTTCCA)

**Sequences of WT enhancers used for Luciferase and Transgenic assays with TAAT and TAAAT motifs highlighted**

a. *CG13222* enhancer

CATAGACCACCAGCCACTGTCAGCACATAACTCAAGCAAACACCTTTGAGGCAGATGCGAAGACAAGTGCTGGAGCAAAAAGACCCAGAATTTGCATATGGGGCTGGCTCATAAACAAGTGCGCGGAAGACGACCGGTGGGCCACCATGAAGATATCAATCAGCCCAGGCCAAAGCCGAAGCCAAACGAAATTCGTTTGTCCTTCTGTCAATGGCCGATAAAAAACCAATAAAGCTGCCAGCCCAGCGGCTTGTTAACACGCAGATAAATTA^*^CACTGGCCGCCCGCGAGATTA^#^CCATCGAGATGCAGTCAGAAGCCGATAGCCAGGGCACAGGAAGCCGCCGATTGTTCTGACCATAACTGAGCTGAGCTTTAGGGCCTGACGAACCGCCGGGCCCGACGATCGAGAACCGCATAAATACGGCCGCCGGTGGGAGCCGCAATTGCAGTCGTGGCACGGCTTTCGATCAGTGACTTCCAGTGGCAGCAACGGATTTGCTTCGGTACGCGATTGA

* M1, M1_A and M1_B – mutations listed in Suppl. Table 1 corresponds to this region.

^#^ M2_A mutation listed in Suppl. Table 1 corresponds to this region.

b. *quadVg* enhancer

AGGAGCTCCCTCCGGAGACCGGGGGCCCAAAAATAGCAACTGCAATTGAGCGGCAGGAGATACCAAAAACTTGCATAGGCTTGCATTTGCGTTGACAACATTCCAAACTCGATTCGAGAGAAAAATATCCAACAACTGGAGAGGAGTTTTGAGGATGCGGACGAGGACGAGGTCGGAGGATGTGGATGTGGATGTGCTTGTGGGGATGTGTAATGTTTGTGCTTGGCTGCCGTCGCGATTCGACAACTTTGGCCGGCACGTTGGCGAGTGTGCCATGCATGCTGATGACGATGAGAATGAGGATGAGGATGAGGATGCGGATGATGATGGTGCTGGTGCGGCTGGGATACTGAATACTGGATACGGGATGCCATGCCGCGTGCCTTTTTTTTCCCGTACCAGAAGCCAGAAGTCGTCATCATCCCATTGCCATTCACTCACTCGCTCAGCTGAGGCCGTGGAATTCCCATTAATGTGCAAACAAGCAGACTGCCAAAGATATTTCCTCTGCAGCTCCTTCAGTTAGCATTTCACTTTCAGCCAGCGGCTTCAAAAGCGAAAGCCGCACTACCTGTCCCCACCTTTCACCTTTTGGCCTAAT^*^GAAGAGAGCGTGGCGATTTAT^#^GACCCGATAACGTTCGATCGCCAGCGTTGACGCATAGTGCGGTCCTGCACAGAGAAAACTATCCTAATCCTCTATGAAGATTCTAAATCAAGTATGGAGTATTCAAACTAAAATCGACTCAACAGCAGTTTTATCTTGCTTAAACTATAGTGAGTTTCAATTAGAAACGACAAAGTACAATCGA

* quad-vg_M2 mutation listed in Suppl. Table 1 corresponds to this region.

\# quad-vg_M and quad-vg_M1 – mutations listed in Suppl. Table 1 corresponds to this region.

c. *ApisVg* enhancer

TTTCTAATCTTCTCGCGAGAAACGAGAGGCTTTCCCCGACTCTTATATAACTGGAAGTTGGACTGCGAGAATGAAAACGGCGCGGCTCTTTTTGCGGCGACGCGTCTTGGATATGTTGGAGCAGAGTCGAGGGAGGGACGGGAGGAGGTGGAGGAGGAGGAGGGAGAGCGAAATGTTTTACGGCCATGCCGTGGCTTTTTGTCGTCGGCATCGATGATTTCGGTGGGAGAACAACGAAGAAAGCACGAGTCCCGAGAGCGTGGCACCGCTATATCGGCCCCCATTAAGCTCTTATTAATT^*^GCGAGCATCTGAGGGGCCGACCGACCTTTCTTGCGCGCTACCTGCGCGCCATTCCGCCTCCGTTCCCTCTCCGCTGCCCGCCGCAAACAAACCCGCTGCACTCCTCGGCCTTCCAACTTTGATGTCGAGTCGATCCTCCTCCAAAGAGATTTATCATGTAAATCGTGGGTAAAAAGCCCGCTCATCCTTCCCTATATCGCGACCCATCGCGTCCTCTTTTTTTCTTTTTTTCTTTTTTTTTTTTCTTCGTGTCCTCGTCACTGTCCACCGCCTTCT

* Apis-vg_M1, Apis-vg_M2, Apis-vg_M3 and Apis-vg_M4 - mutations listed in Suppl. Table 1 corresponds to this region.

**Table S1. Details of mutant constructs and Primer sequences**

| **Name of construct** | **Description** | **Vector backbone** | **Primer sequence (Forward)** |
| --- | --- | --- | --- |
| ***CG13222* enhancer constructs (For Luciferase Assay and EMSA)** | | | |
| M1_A | Only TAAAT motif mutated at site1 | pGl3-DE5 | GCTTGTTAACACGCAGAGGGATTACACTGGCCGCCCGCGAGATT |
| M1_B  (only used for EMSA) | Only TAAT motif mutated at site1. This was used for EMSA only |  | CGCAGATAAATGGCACTGGCC |
| M2_A | TAAAT motif introduced in place of TAAT motif in site2 | pGl3-DE5 | CACTGGCCGCCCGCGAGATTTACCATCGAGATGCAGTCAG |
| ***CG13222* enhancer constructs (For *Drosophila* Transgenics)** | | | |
| M1 | Both TAAT and TAAAT motifs mutated (Hersh et. al 2007) | pH-Stinger-attb | GCTTGTTAACACGCAGATAACGCGCACTGGCCGCCCGCGAGATT |
| M1_A | Only TAAAT motif mutated at site1 | pH-Stinger-attb | GCTTGTTAACACGCAGAGGGATTACACTGGCCGCCCGCGAGATT |
| M2_A | TAAAT motif introduced in place of TAAT motif in site2 | pH-Stinger-attb | CACTGGCCGCCCGCGAGATTTACCATCGAGATGCAGTCAG |
| ***quad-vg* enhancer constructs** | | | |
| *quad-vg*_M | TAAAT motif mutated | pH-Stinger-attb | GATCGAACGTTATCGGGTCCGCGCCGCCACGCTCTCTTCATTAG |
| *quad-vg*_M1 | TAAAT motif mutated to TAAT motif | pH-Stinger-attb | GATCGAACGTTATCGGGTCTAATCGCCACGCTCTCTTCATTAG |
| *quad-vg*_M2 | TAAT motif mutated | pH-Stinger-attb | TCGCCACGCTCTCTTCGCGCGGCCAAAAGGTGAAAG |
| ***Apis-vg* enhancer constructs** | | | |
| *Apis-vg*_M1 | Mutation replacing the entire TAAT motif with TAAAT motif | pH-Stinger-attb | TCGGCCCCCATTAAGCTCTTCGATTTATGAGCGAGCATCTGAGGGGCCGA |
| *Apis-vg*_M2 | Mutation replacing the TAAT motif with a 25bp cassette from *quad-vg* | pH-Stinger-attb | TCGGCCCCCATTAAGCTCTTGAGAGCGTGGCGATTTATGAGCGAGCATCTGAGGGGCCGA |
| *Apis-vg*_M3 | Mutation replacing the TAAT motif with par of the cassette from quad-vg (TAAAT site present) | pH-Stinger-attb | TCGGCCCCCATTAAGCTCTTTAATGAAGAGAGCGTGGCGATTTATGAGCGAGCATCTGAGGGGCCGA |
| *Apis-vg*_M4 | Mutation replacing the TAAT motif with part of the cassette from quad-vg (TAAAT site absent) | pH-Stinger-attb | TCGGCCCCCATTAAGCTCTTTAATGAAGAGAGGCGAGCATCTGAGGGGCCGA |

**Figure S1**

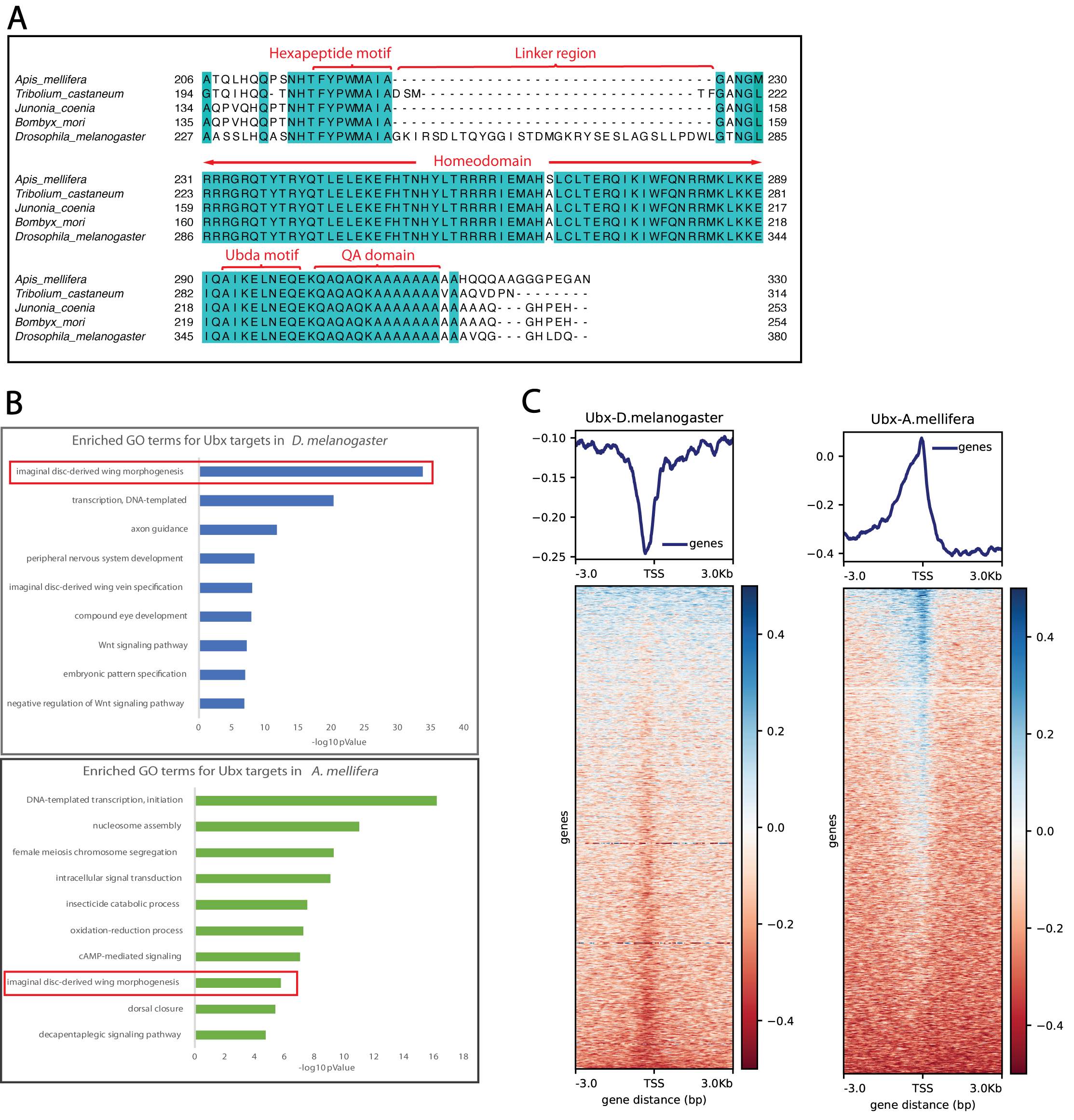

**Legend Figure S1**

**A)** Protein sequence comparison of Ubx from different insect species showing conservation in the homeodomain region as well as protein interacting motifs (HX, Ubda). The linker region, however, is variable with Dipterans like *Drosophila* displaying a longer linker sequence as compared to others.

**B)** Gene ontology analyses of direct targets of Ubx indicate common wing patterning genes targeted by Ubx in both *Apis* hindwings and *Drosophila* halteres. However, the proportion of wing patterning targets in *Drosophila* is larger as compared to *Apis.*

**C)** Heatmap analysis indicate clustering of Ubx-binding sites away from the transcriptional start sites (TSS) in *Drosophila*, while in *Apis* they are very proximal to TSS.

**Table S1**

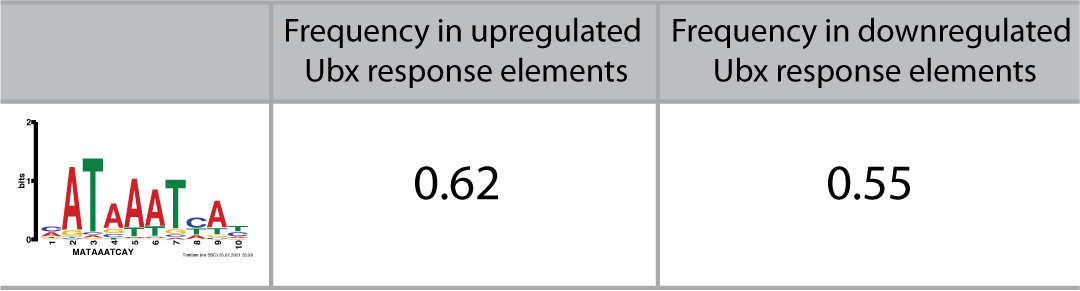

**Legend Table S1:** Frequency comparison of the TAAAT motif in Ubx response elements that correspond to genes that are upregulated and downregulated in the haltere. The frequency of the TAAAT motif is slightly higher in the upregulated category (1.4 folds) as compared to the downregulated category

**Figure S2**

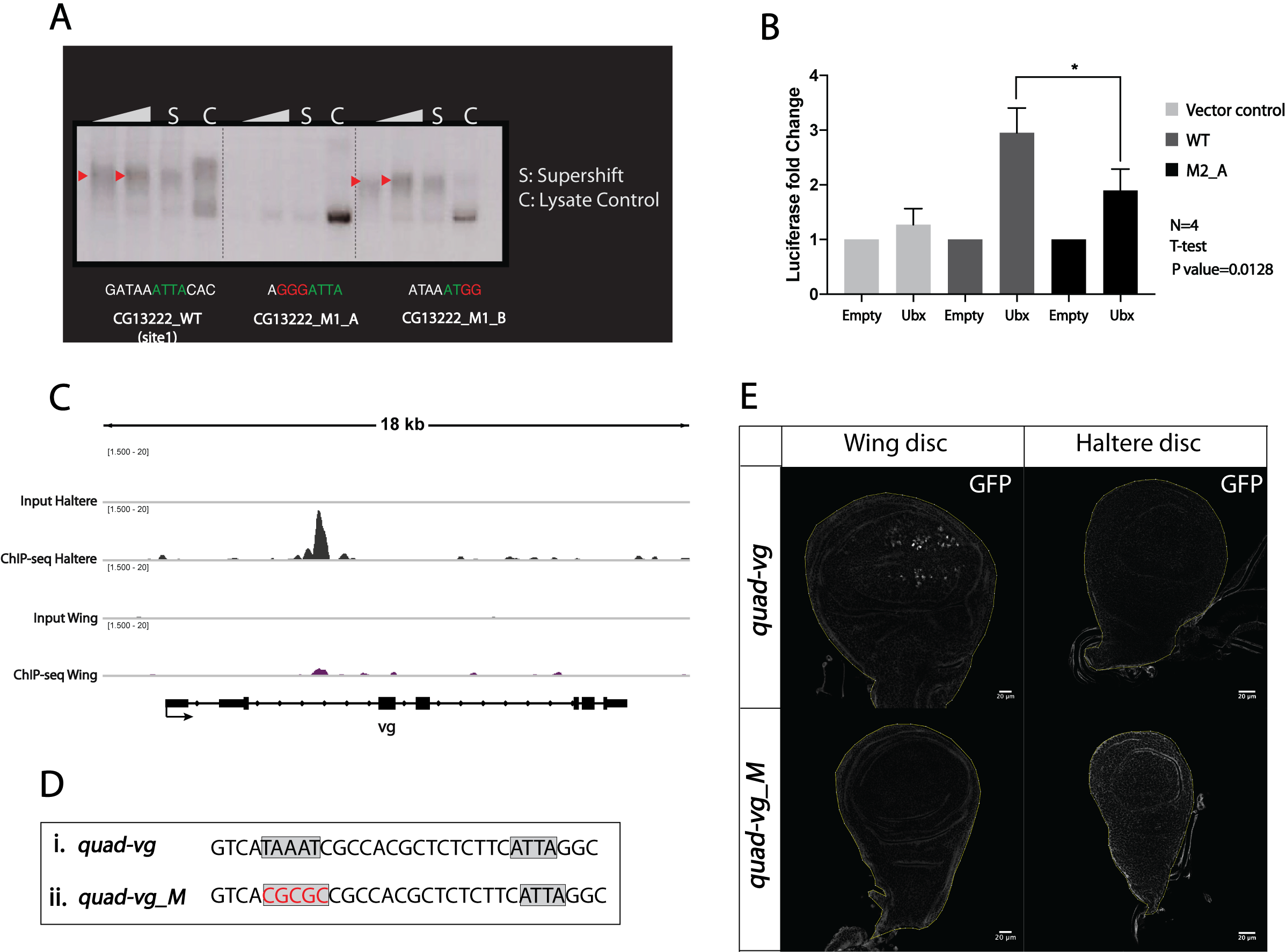

**Legend Figure S2:**

**A)** EMSA showing the difference of Ubx binding to the CG13222 enhancer in presence and absence of the TAAAT site.

**B)** Mutation of the TAAT site to TAAAT in site2 of the *CG13222* enhancer leads to significant difference in reporter expression driven by the mutant enhancer. For statistical analysis, t-test was performed (two-tailed).

**C)** Ubx peak observed in the intronic region of the *vg* gene.

**D)** Sequence of the region of quad-vg enhancer containing the TAAT and TAAAT motifs. Mutations were generated to alter the TAAAT motif (quad-vg_M)

**E)** Expression driven in the wing and haltere imaginal discs by the *quad-vg* and its mutant enhancer. Mutations in the TAAAT motif lead to inactivation of the enhancer element and loss of expression from the wing imaginal discs as well.

**Figure S3**

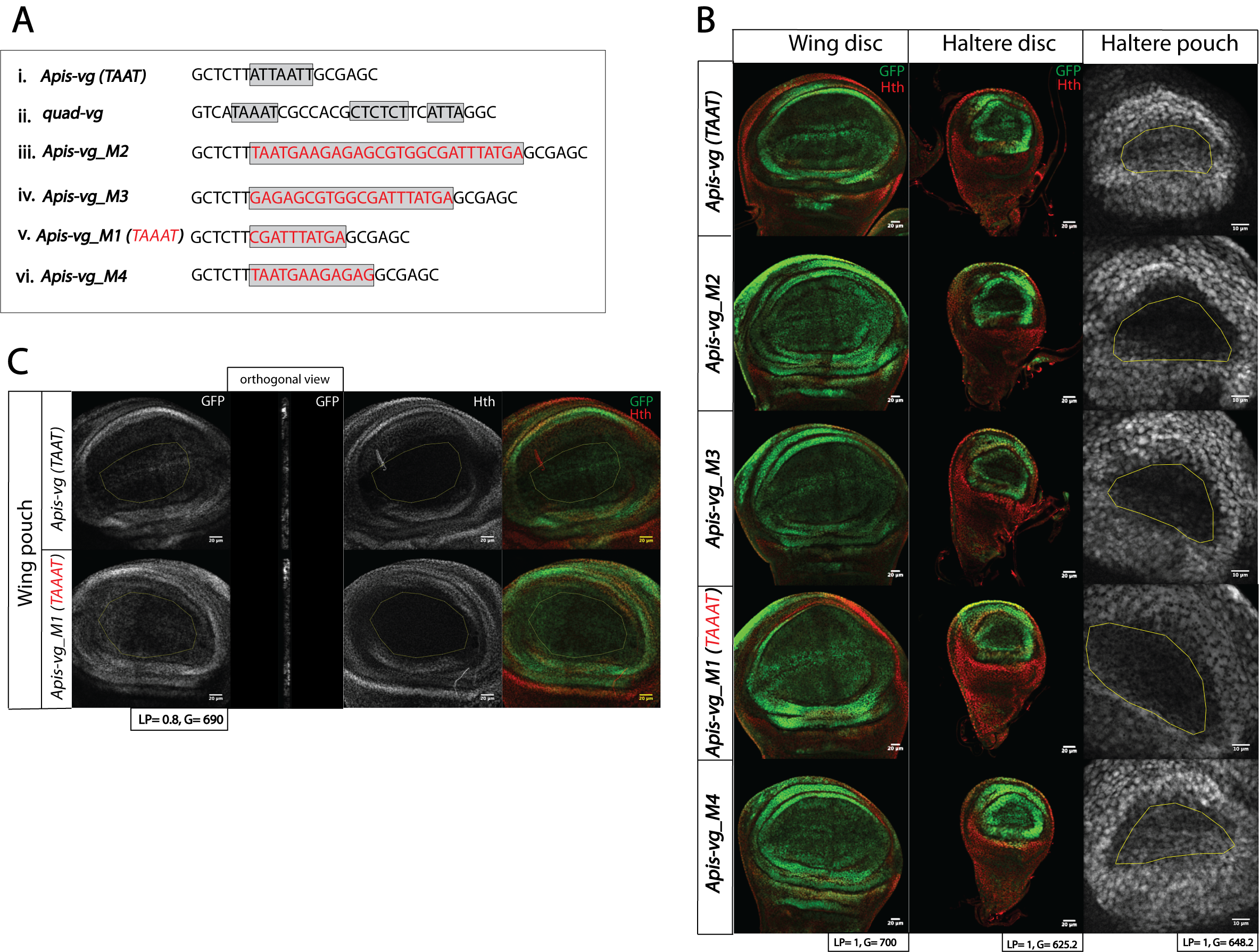

**Legend Figure S3:**

**A)** Sequence of i) part of the 575bp enhancer of the *vestigial* gene in *Apis (Apis-vg)*, showing the presence of a TAAT motif and (ii) part of 806bp enhancer of the *vestigial* gene in *Drosophila (quad-vg)* containing the TAAT and TAAAT motifs. The *quad-vg* also contains a GAGA factor binding site between the TAAT and TAAAT motifs. (iii-vi) Various mutant versions of *Apis-vg* to bring in features of *quad-vg* of *Drosophila*.

**B)** Expression driven by the *Apis-vg* and its mutant enhancers in third instar wing and haltere imaginal discs. Loss of expression in the haltere pouch is correlated to the presence of TAAAT motif not the TAAT motif.

**C)** Magnified images of the wing pouch of *Drosophila* transgenics expressing GFP as driven by *Apis-vg* and *Apis-vg_M1* enhancers. Please note both the wildtype and the mutant versions of the enhancer drive similar levels and patterns of GFP expression.
